# Non-invasive monitoring of microbiota and host metabolism using Secondary electrospray ionization-Mass spectrometry

**DOI:** 10.1101/2022.05.25.493434

**Authors:** Jiayi Lan, Giorgia Greter, Bettina Streckenbach, Markus Arnoldini, Renato Zenobi, Emma Slack

## Abstract

The metabolic “handshake” between the microbiota and its mammalian host is a complex, dynamic process with potentially major influences on health. Dissecting the interaction between microbial species/strains and metabolites found in host tissues has been a challenge due to the high diversity of a complete microbiota and the requirement for invasive sampling, which precludes high-resolution longitudinal analysis. Here we demonstrate that secondary electrospray ionization mass spectrometry can be used to non-invasively monitor metabolic activity of the intestinal microbiome of a live, awake mouse. This was achieved via analysis of the headspace volatile and semi-volatile metabolome of individual gut microbiota bacterial species growing in pure culture, as well as from live gnotobiotic mice specifically colonized with these microbes (i.e. metabolites released to the atmosphere via breath, the skin and from the gut). The microbial origin of these compounds was confirmed by feeding of heavy-isotope labeled microbiota-accessible sugars. This reveals that the microbiota is a major contributor to the released metabolites of a whole live mouse, and that it is possible to capture the catabolism of sugars and cross-feeding within the gut microbiota of a living animal using volatile/semi-volatile metabolite monitoring.

## 1. Introduction

The gastrointestinal tract is colonized by a complex ecosystem of microbes, referred to as the gut microbiota[1], which plays a major role in modulating host metabolism and immune function. Exchange of metabolites between the microbes and their host is an integral mechanism in this interaction [2, 3]. Correspondingly, our ability to measure the production of small molecules produced by a host and its microbiota has the potential to advance our understanding of this highly complex relationship [4, 5]. To date, most of these efforts have focused on quantification of metabolites in body fluids, often requiring invasive sampling. Moreover, the easily accessible fecal metabolites are often used as a proxy for microbial metabolism that is most active in the upper large intestine. Acquiring meaningful, time-resolved data on microbial activity in live animals is thus challenging. However, this can be done by monitoring small molecule metabolites released by the experimental animals to the gas phase, including volatile, semi-volatile and low volatility species (hereafter, for simplicity, referred to as the “volatilome”). Progress in ambient mass spectrometry (MS) now renders it possible to non-invasively monitor the volatilome in breath [6, 7, 8] and exhaust gases [9, 10]. Notably this includes the ability to detect short-chain fatty acids (SCFA), which are of known interest in host-microbe interactions. SCFA have already been extensively studied, for example in epithelial gene expression and metabolism [11]. This suggested the possibility to apply ambient mass spectrometry technology to the study of microbiome function.

Studies on plasma have shown microbiota-dependent effects on a variety of host metabolites, including on amino acids [3, 12], fatty acids [13, 14], lipid, bile acids [15] and xenobiotics [16]. When conventionally colonized mice are compared to germ-free (GF) mice, 10 % of identified features in common between these mice differ in signal intensity [17]. Furthermore, it was previously suggested that plasma metabolites in the host are predictive of human gut microbiota diversity, and metabolites in the host can be used as predictors of microbial phyla abundance [18].

Mass-spectrometry based approaches, including Matrix-Assisted Laser Desorption-MS (MALDI-MS) [19], Liquid Chromatography-MS (LC-MS) [20, 21] and Gas Chromatography-MS (GC-MS) [22] have been applied to microbiome research to profile the metabolome of various bacterial or host-tissue samples at specific time points. Time-resolved monitoring of host metabolism is possible using sensors measuring light gases released by the organism (e.g., H_2_, CH_4_, O_2_ and CO_2_, [13, 23, 24, 25]). This data allows monitoring of some aspects of host and microbiota metabolism, such as host energy expenditure, respiratory quotient and microbial hydrogen production. However, due to the limit of available sensors, only a small number of simple gases or volatiles can currently be studied using these methods.

Recent advances in ambient mass spectrometry made it possible to detect complex volatile and semi-volatile metabolites in the gas phase. Secondary electrospray ionization mass spectrometry (SESI-MS) which has been developed in the past decades [26], is compatible with high-resolution MS, and has a relatively fast response and broad detection range [27, 28] when compared to conventional MS-based detection methods for volatiles. For example, SESI-MS allows direct detection without pre-concentration and separation (compared to GC-MS [29]), it can detect volatiles ionized in both positive and negative mode (compared to proton transfer reaction mass spectrometry [30], which only allows positive ion mode) and it can be easily mounted on a high resolution MS (compared to selected ion flow tube mass spectrometry mass spectrometry, [31]). SESI-MS al-lows detection of molecules in the gas phase down to sub-ppt concentrations with minimum detectable vapor pressure down to 10^*−*7^ bar [32, 33]. It is therefore the method of choice for efficient, real-time detection of volatile metabolites in biological systems. Previously, SESI-MS has been used for identifying biomarkers for various lung pathogens in human breath [34], and to capture the metabolic responses of *Staphylococcus aureus* to antibiotic treatment in the headspace of microbial batch cultures [35]. However, due to the high complexity of the gut microbial community, it is usually difficult to assign metabolic signals to a specific origin and to decipher causal relations between changes in bacterial and host metabolism, thus the application of SESI-MS in gut microbiota research is yet to be explored.

Here, we present a SESI-MS based approach to simultaneously study microbiome and host metabolism, by monitoring volatilome released from live animals in which we can manipulate the microbial composition: ranging from germ-free, via a simplified 3-species microbiota to a complete “specific-pathogen-free” (SPF) microbiota. By first identifying volatile markers for each member of the reduced microbiota, we were able to track bacterial metabolism inside a host non-invasively, while at the same time monitoring the changes in host metabolism. Feeding of microbiota-available heavy isotope-labeled sugars allowed us to unambiguously identify the microbial origin of volatilome, and to directly observe microbial cross-feeding in live animals.

## 2. Results

### 2.1. A setup to identify small-molecule metabolites in gas released by animals and their microbes

Invasive interventions are a major stress for research animals and thereby influence the results of any metabolome analysis. We therefore developed a SESI-MS-based approach to measure the volatilome released to the local atmosphere by unperturbed animals and their microbiota (Fig 1A). To measure the volatilome of the murine host alone, germ-free mice were placed in a red plastic tunnel (7 cm diameter, 11 cm length) fed with sterile medical air, which could be directly connected to the SESI-MS inlet for measurements (Fig 1B, D). In order to benchmark the system for microbial metabolite analysis, and to identify metabolites characteristic for particular bacterial species, we also established a custom-made headspace sampling system coupled to the SESI-MS (headspace-SESI, Fig 1B, C), which makes it possible to identify metabolic signatures of microbial cultures. By combining the volatilome of germ-free mice and the headspace volatilome of individual cultures, we established identifiers for the different parts of the host-microbiota system, i.e., metabolic signatures for different microbiota members and baseline metabolic profile of the uncolonized host. In a next step, by measuring the volatilome of colonized mice, we can infer the effect of metabolic interactions between the host and its microbiota.

**Figure 1:**
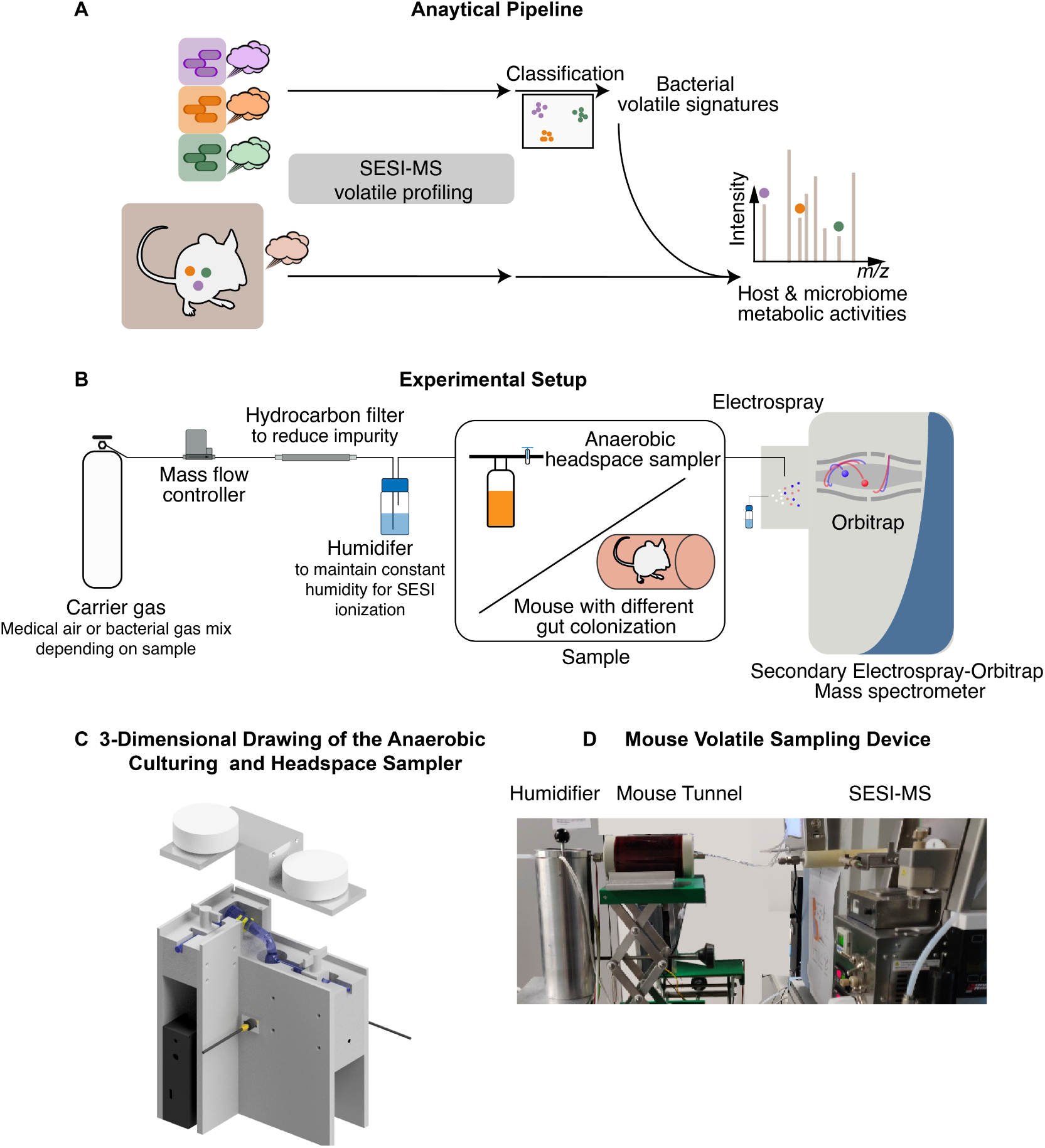
Workflow and experimental settings for non-invasive monitoring of microbiome and host metabolism. **A**, Schematic workflow for non-invasive monitoring of host-microbiome metabolism. **B**, Experimental setups for SESI-MS measurement of either bacterial cultures or mice. **C**, 3-dimensional drawing of the anaerobic culturing and headspace sampler. The sampler consists of a set of glass containers (1.5 cm outer-diameter quartz cylinder flask with gas valve on both inlet and outlet), a home-built optical density meter and a metal holder kept at 37 °C. **D**, Picture of experimental setup for measuring mouse volatiles. The mouse was kept in an air-tight red plastic tunnel in order to reduce its stress level. Humidified medical air at a flow rate of 0.3 L/min constantly flows through the tunnel and transfers the released volatiles into SESI-MS.

### 2.2. Distinction of different colonization states based on the volatilome

As a first proof of principle, we tested whether it is possible to distinguish different colonization states of mice using our SESI-MS setup. We measured the volatilome of C57BL/6 mice that were either germ-free, or colonized with two different microbiota. The simpler microbiota (Easily Accessible Microbiota, EAM) consists of three bacterial strains, *Bacteroides thetaiotaomicron, Escherichia coli*, and *Eubacterium rectale*. These are common members of the typical human gut microbiota, representing the three most abundant bacterial phyla. The three microbes colonize the mouse large intestine to a density of between 10^9^ and 10^11^ bacteria per gram of gut content (Fig 2C, D). To represent a more complex microbiota we used conventional specific pathogen-free (SPF) mice raised in the EPIC mouse facility at ETH Zurich. Using the SESI-MS setup depicted in Fig 1D, we measured the volatilome released by these three different groups of mice for 500 seconds (around 8 minutes). Principal component analysis could clearly distinguish the three different colonization states based on the detected metabolite pool (Fig 2C). We then looked for bacterial signatures in the data, focusing initially on the SCFAs acetate, propionate, and butyrate as markers for bacterial metabolism. These are typical end-products of microbial fermentation in the intestine and are known to be correlated with microbial activity [36, 37, 38]. As expected, we saw increased levels of all three SCFAs in colonized mice as compared to GF mice, with butyrate detected at higher levels in the 3-species microbiota-colonized mice, whereas acetate and propionate were higher in the SPF microbiota (Fig 2B).

**Figure 2:**
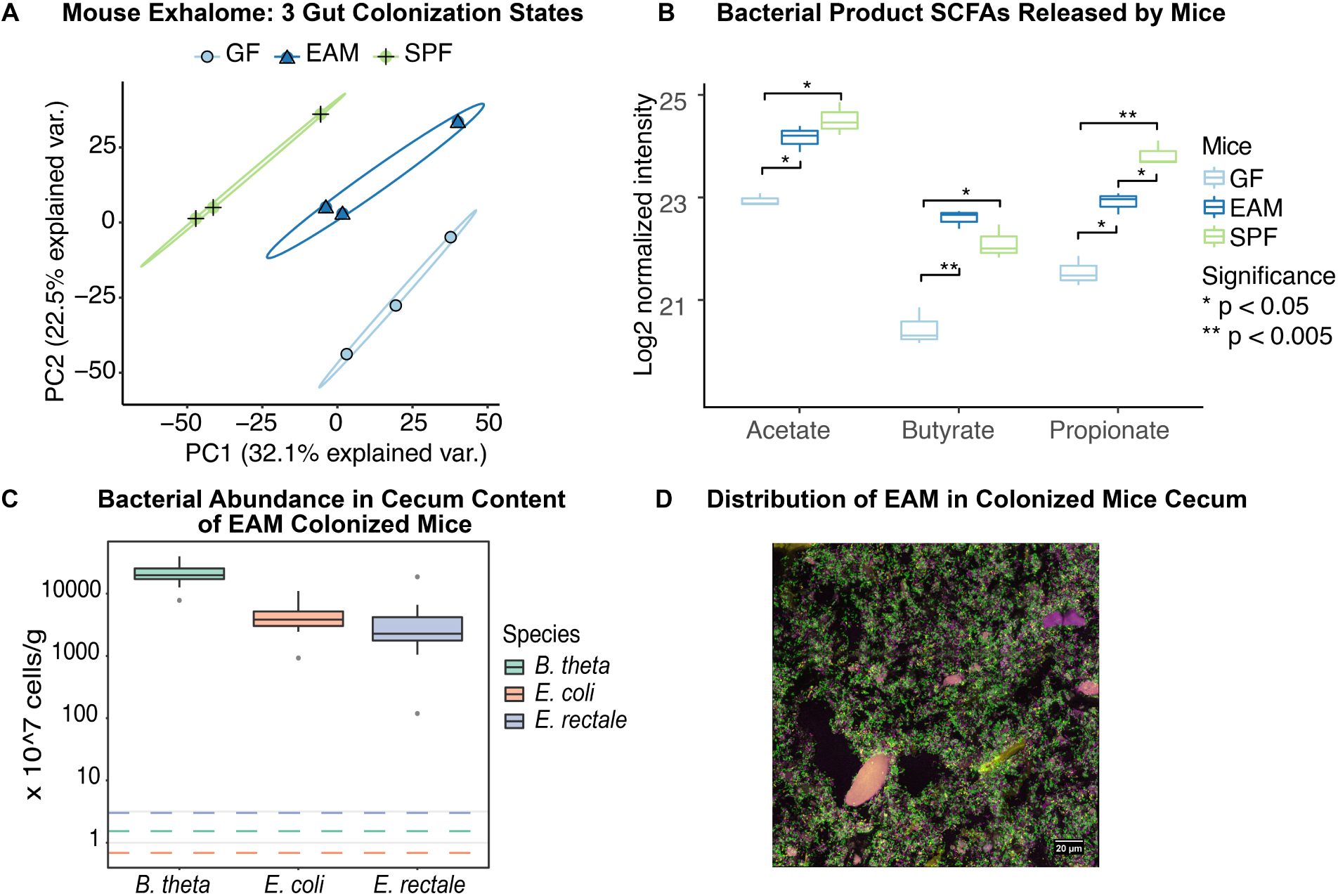
Different gut colonization can be detected in emitted metabolites of the host. **A**, PCA plot of volatilome of mice with different gut microbiota: germ-free (GF), EAM-colonized (EAM) and SPF. **B**, Levels of typical bacterial fermentation products, SCFAs, released by differently colonized mice. **C**, bacterial abundance in cecum content of EAM colonized mice, quantified using qPCR. **D**, Fluorescence *in situ* hybridization (FISH) imaging of EAM-colonized mice gut showing the distribution of EAM in colonized mice cecum.

### 2.3. Disentangling host and microbiota signals in a minimal microbiota

After establishing that our experimental setup can distinguish between different colonization states and can identify signature microbial metabolites in the volatilome around a live, unperturbed mouse, we next aimed to delineate the contribution of the different parts of the host-microbiota system. For this, we used the more tractable 3-member microbiota (EAM) as this allowed us to compare the pure-culture metabolome to the colonized mice. Headspace-SESI-MS metabolome profiles of the different strains during stationary growth, as well as of the anaerobic culture medium alone clearly clustered together repeats of the same bacterial strain and separated the strains/media from each other (Fig 3A). As gut bacteria are typically growth limited by nutrient availability, late-log/early stationary phase is a reasonable approximation of the majority of cells [39]. Discriminative features for each of the bacterial strains were identified using SparseSVM models (one-vs-all, [40, 41]). We validated the model training on separately acquired test data, which yielded model accuracies of 1.00, 0.89, and 0.96 for *B. theta, E. coli*, and *E. rectale*, respectively. The discriminative mass/charge (*m/z*) features identified in this process were then annotated with various approaches, including online MS2 fragmentation and matching with METLIN [42] or MassBank [43] database, and accurate mass matching with the KEGG database ([44]) as a reference (organism codes: bth for *B. theta*, ecx for *E. coli*, ere for *E. rectale*). Putative annotation can be found in Tab. S1. Acetoin, isopentyl acetate and valeric acid were found to be characteristic for *B. theta*. Hydroxy butyric acid, p-cresol, caproic acid and indole were found to be characteristics for *E. rectale*. Catechol, L-aspartic acid, shikimic acid, pyridoxal and pyridoxine were found to be characteristic for *E. coli* (Tab. S1).

**Figure 3:**
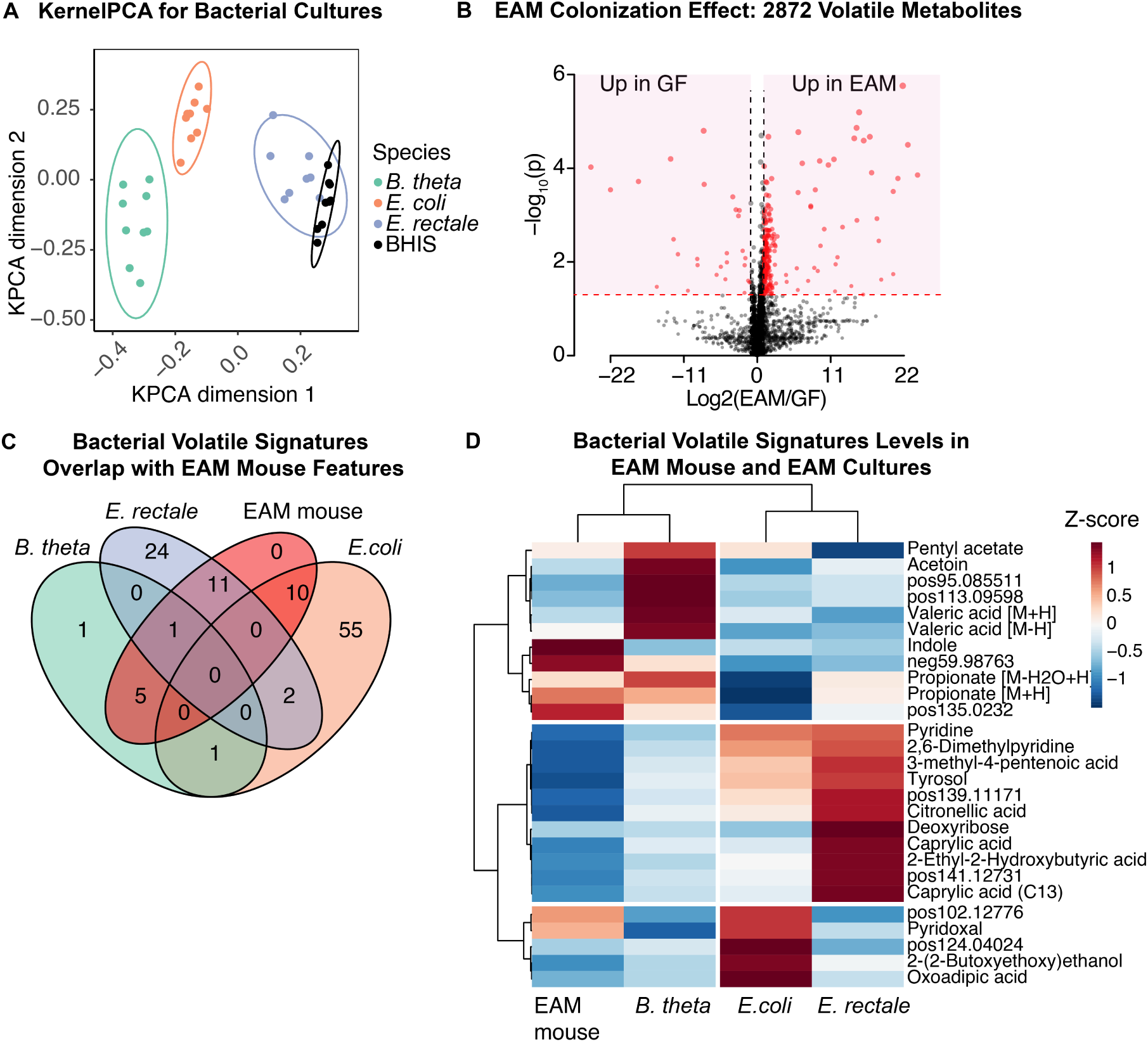
Tracking of bacterial volatile signatures detected *in vitro* in EAM-colonized mice. **A**, KernelPCA plot of the *in vitro* EAM species and the BHIS culture media. **B**, Volcano plot comparing volatilome of GF mice and EAM mice (41 FDR adj. p *<* 0.05 & fold change *>* 2, 253 raw p *<* 0.05 & fold change *>* 2). **C**, Overlaps between bacterial volatile signatures identified *in vitro* and EAM mouse volatile features. **D**, Hierarchical clustering showing normalized intensities of bacterial volatile signatures in culture and in EAM-colonized mouse (volatile signatures’ intensity in EAM mice subtracted by intensity in GF mice).

Next, we compared how the volatilome differs between GF mice and mice colonized with the three-member microbiota, by analyzing the data depicted in Fig 2A in more detail. Out of 2872 detected features, 41 were significantly altered between the GF and the colonized mice (FDR adjusted p value *<* 0.05 & fold changes *>* 2, Fig 3B). The differences clustered in the pathways for butyrate/propionate metabolism and the pathways for metabolism of various amino acids (Tab. S2). These results are consistent with previous results using different measurement methods, namely NMR analysis of mice colonocytes showing increased SCFA in colonized animals [45], and bacterial activities modulating host amino acid metabolism in LC-MS based analysis in mice hepatic portal vein blood [3].

It has been shown recently that signature metabolites for microbial strains in culture can also be detected if those strains colonize a host [4]. We therefore tested whether the set of discriminative features we have identified in microbial cultures (Fig 3A) can also be identified in the volatilome of a colonized mouse. We subtracted the volatilome profiles of GF mice from those of colonized mice to correct for the host influence, and compared the resulting profiles to that obtained from microbial cultures. Of all 110 selected features that discriminated between bacterial strains in pure culture, 27 were detected in the colonized mouse dataset (Fig 3C). As in Fig 3D, a hierarchical clustering of the identified features shows that the features of the EAM colonized mice cluster most closely together with the *B. theta* features, which is in line with the observation that *B. theta* colonizes with a roughly 10-fold higher abundance than the other two bacterial strains (Fig 2C).

### 2.4. Monitoring a microbiota-targeted intervention non-invasively

Despite the identification of microbe-associated metabolites in the volatilome of colonized mice, it remains possible that some of these metabolites could be the result of microbiome-driven alterations in host metabolic activity. To categorically demonstrate that the metabolites we detect using our system are indeed directly reflective of microbial metabolism in the host, we fed fasted mice ^13^C-labeled D-arabinose (^13^C-ara, Fig 4A). This sugar is not metabolized by the mouse but can be utilized by *B. theta* in our EAM model as well as by other microbes in complete microbiota (Fig S4B, [46, 47]) which metabolize it into heavy-isotope labeled metabolites that can be detected using SESI-MS (Fig 4B). When comparing the total volatilome profiles from ^13^C-ara fed mice with those of the control groups, we found an enrichment of metabolic features in various pathways, namely amino acid metabolism, butyrate metabolism, fatty acid biosynthesis and central metabolism (TCA cycle) (Table S3). Focusing on 27 of the unlabeled microbial features that were identified in both the headspace SESI-MS (Fig 3C) and the volatilome of EAM colonized mice (Fig 4C), we could see changes in four of these features after ^13^C-ara feeding. Intriguingly, three of those are *E. coli* specific features, whereas one is *B. theta* specific. Our *E. coli* strain cannot utilize D-arabinose directly (Fig S4B), which indicated that there may be indirect effects of D-arabinose on *E. coli* metabolism.

**Figure 4:**
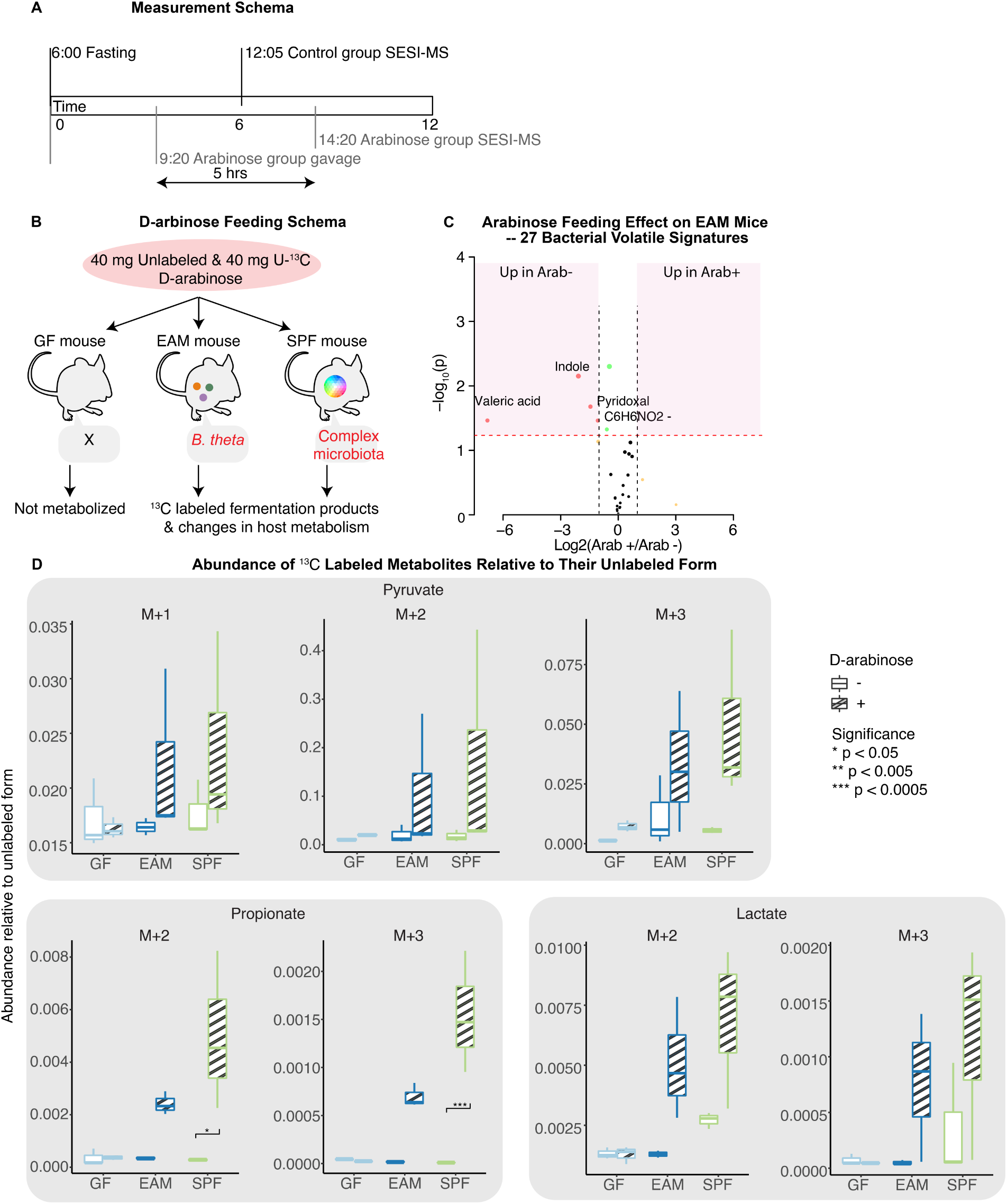
Bacteria-specific isotope labeled carbon source identifies direct microbial metabolites in the volatile metabolome^32^. **A**, Measurement schema of the control group and the arabinose group. **B**, D-arabinose feeding and uptake schema for mice with various gut colonization. **C**, Volcano plot showing arabinose feeding effect on bacterial volatile signatures detected in EAM mice. (4 raw p *<* 0.05 & fold change *>* 2) **D**, Labelling patterns of bacteria-related metabolites, including pyruvate, propionate and lactate.

When considering the ^13^C-labeled components of the volatilome, we could clearly observe that SPF and EAM mice, but not GF mice, incorporated D-arabinose-derived carbon into metabolites such as pyruvate, propionate and lactate. This both confirmed the inert behavior of ^13^C-ara in the absence of microbial metabolism, and the potential of SESI-MS to directly monitor microbial metabolism within a live unperturbed host (Fig 4D).

### 2.5. Detection of cross-feeding between microbiota members inside the host gut using SESI-MS

In the EAM host-microbiota system, the only species competent for growth on D-arabinose is *B. theta* (Fig 4B, Fig S4B). After feeding the EAM mice ^13^Cara, all metabolites with increased ^13^C labels therefore directly or indirectly result from *B. theta* breaking down this molecule. Intriguingly, in addition to metabolites that are well known intermediates or products of metabolic pathways found in *B. theta*, such as pyruvate and propionate (Fig 4D), we found an increase in ^13^C labeled versions of butyrate. Butyrate is a typical metabolite produced by *E. rectale* [48] (Fig 5A) in large quantities, but cannot be produced directly by *B. theta*. This is consistent with the observation that *E. rectale* can take up acetate produced by other microbes, which can then be incorporated into butyrate [49, 50](Fig 5B). We observed an increase in acetate carrying two ^13^C labels (M+2 acetate) in the metabolite pool, and increases in double (M+2) and quadruple (M+4) ^13^C labeled butyrate, respectively. The simplest explanation of this data is that M+2 acetate produced by *B. theta* is being cross-fed into *E. rectale*, where it is then incorporated into butyrate: i.e. we can use SESI-MS to quantify cross-feeding *in vivo*.

**Figure 5:**
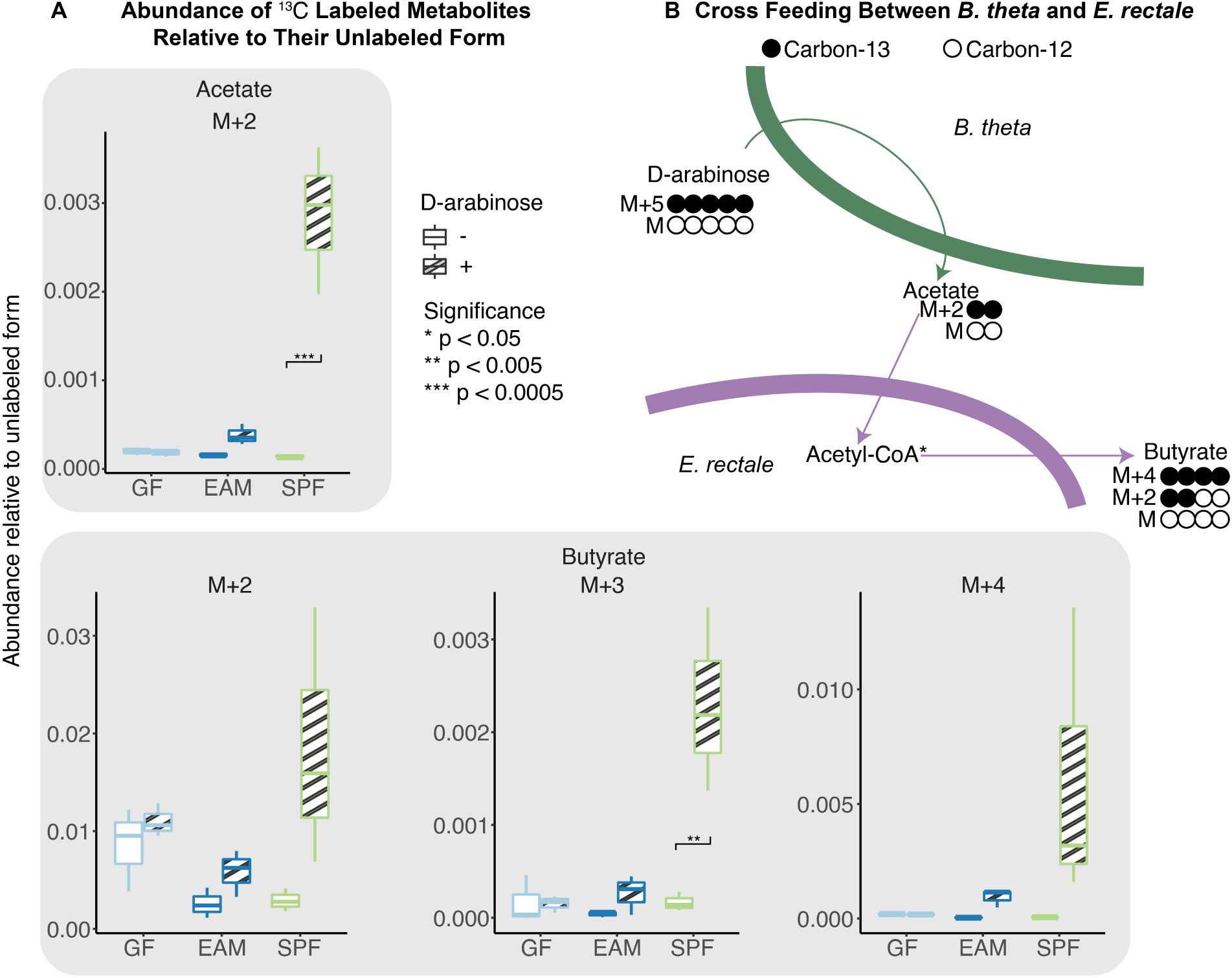
Bacteria-specific isotope labeled carbon source reveals crossfeeding *in vivo*. **A**, Labelling patterns of acetate and butyrate released by mice with different microbiota. **A**, Generating of differently labeled butyrate through cross feeding between *B. theta* and *E. rectale*. Black circles indicate Carbon-13 and white circles indicate Carbon-12.

## 3. Discussion

In this study, we have shown that SESI-MS of the mouse volatilome is a powerful, non-invasive tool to analyze host-microbiota interactions. By comparing the SESI-MS signatures of the host-microbiota system in gnotobiotic animals with those of the colonizing microbes in isolation, we were able to identify the metabolite footprint of the most abundant microbes (Fig 3D). By feeding with isotope labeled sugars that can only be used by the microbes and not by the host, we were able to conclusively show the presence of microbial metabolites in the volatilome, and could use these metabolites to demonstrate within-host cross-feeding between intestinal bacteria.

It should be noted that SESI-MS analysis of air around an animal clearly does not, and is not expected to, detect all produced metabolites. Very small metabolites, such as H_2_ and CO_2_, have too low a molecular weight to be measured by our mass spectrometer, and high molecular weight metabolites present in the gas phase with extremely low concentration fall below the detection limit of SESI-MS. However, within the accessible molecular weight window for observing host-microbiota metabolism, we were able to recapitulate metabolic changes upon microbial colonization of a GF mouse which have previously been measured in host tissues and plasma (e.g. amino acid metabolism, xenobiotic metabolism, Tab S2, [13, 51]). Given that SESI-MS is non-invasive and does not preclude the later analysis of e.g. host tissue, and that many metabolic pathways have profound circadian rhythms, a potential application is therefore the identification of the optimal time-points for endpoint analysis [9].

We envisage one of the most powerful features of the SESI-MS approach to be the possibility of collecting time-resolved data on metabolic processes of a host-microbiota system in an undisturbed way. This has, up to now, only been possible by collecting fecal samples, which comes with a set of limitations. First, fecal samples are not representative of active microbial metabolism [52], which happens largely in the upper part of the large intestine [53, 54, 55]. Second, repeated fecal sampling, even though minimally invasive, will disturb an animal’s routine if conducted over longer time periods and will therefore affect its behavior and potentially the experimental outcomes. And third, sampling of fecal material is limited in its time resolution, which can lead to gaps in information when measuring the effects of experimental interventions that work with a time delay, which is true for most experimental interventions with an effect inside the host. The SESI-MS approach presented here is not affected by any of these issues.

Given availability of the equipment, the workflow as presented here is relatively easy to implement and allows room for customization. SESI sources are commercially available and can be interfaced with commercial high-resolution mass spectrometers [27, 28, 56]. The data generated from SESI-MS is compatible with standard software for direct-injection MS-based metabolomics [57, 58], and online MS2 fragmentation for compound identification can be performed [59, 60]. Development is still needed for collecting and processing time-series data, since most current MS-based metabolomics datasets have low time-resolution or do not consider time-dependent changes at all. One example where time series of SESI-MS data at a minute-level resolution has been successfully used was for continuous monitoring of human metabolism during sleep [57].

In conclusion, we describe a new method for tracking metabolic changes in host-microbiota systems non-invasively. This method has the potential to generate metabolomic data for such systems at an unprecedented time resolution, and with minimal to no system perturbation. Applying this system to gnotobiotic and germ-free mice has considerable potential to reveal relevant microbiota functions and to improve our understanding and diagnosis of microbiota-associated conditions.

## 4. Methods

### 4.1. Animal experiments

Wild type C57BL/6 mice were maintained on a standard diet at the ETH Phenomics Center. Mice were re-derived Specific Pathogen Free (SPF) or rederived germ-free and bred in individually ventilated cages under strict hygienic conditions. Gnotobiotic mice were colonized with EAM strains (the Easily Accessible Microbiota, see ‘Bacterial strains and media’) via oral gavage of mixed overnight cultured bacteria in hemin-supplemented (0.25 mg/*μ*L) brain heart infusion (BHIS) medium. All experiments began between 8-9 weeks of age. All animal experiments were approved by the Swiss Kantonal authorities (License ZH120/19 and ZH058/19; Kantonales Veterinäramt Zürich) and performed according to the legal and ethical requirements.

### 4.2. Bacterial strains and media

For preparing bacterial cultures, *Escherichia coli* O9 HS, *Bacteroides thetaiotaomicron* VPI-5482 and *Eubacterium rectale* ATCC 33656 was overnight cultured and diluted to OD = 0.5 before measured by HS-SESI. For identifying volatile signatures of the three bacterial strains, hemin-supplemented (0.25 mg/*μ*L) brain heart infusion broth was used as growth medium, 10 % CO_2_ in nitrogen was used as atmospheric and carrier gas for HS-SESI measurement. For assessing anaerobic growth of the EAM strains inside of the headspace sampler, overnight cultures of EAM strains were subcultured to OD 0.05 in hemin-supplemented brain heart infusion broth and cultured with 10% CO_2_ and N_2_ as atmospheric and carrier gas for HS-SESI measurement.

For measuring bacterial growth in D-Arabinose, overnight cultures of EAM strains were diluted 1:1000 in D-arabinose-supplemented (0.02 M) epsilon minimal media (1 % tryptone, 0.02 M NH_4_Cl, 50 mM NaCl, 50 *μ*M MnCl_2_, 50 *μ*M CoCl_2_, 50 *μ*M MgCl_2_, 50 *μ*M CaCl_2_, 4 *μ*M FeSO_4_, 20 mM NaHCO_3_, 5 mM cysteine, 1.2 mg/L hemin, 1 mg/L menadione, 2 mg/L folinic acid, 1 mg/L B12, buffered to pH 7) and cultured for 8 hours. Cultures were subcultured to OD 0.05 in 500 *μ*L epsilon medium in a flat-bottom 24-well plate (Techno Plastic Products, Switzerland) and bacterial growth was measured using a Infinite 200 Pro plate reader (Tecan Group Ltd., Switzerland).

### 4.3. EAM quantification

Fourteen 8-9 week old mixed-sex EAM-colonized mice were kept under sterile conditions in two experimental isolators and kept on an *ad libitum* diet for 4 weeks. After euthanasia, cecum tip was removed and used for fluorescence *in situ* hybridization (FISH). Cecum content was collected and bacterial DNA was extracted using the DNeasy PowerSoil Pro Kit (Qiagen, Germany). qPCR was performed using the FastStart Universal SYBR Green Master Mix (Roche, Switzerland). Primers (Supplementary Tab. S4) were diluted to a final concentration of 1 *μ*M. Bacteria were quantified using a standard curve of known bacterial DNA concentration. DNA was amplified using a QuantStudio 7 Flex instrument (Applied Biosystems, MA, USA) with an initial denaturation step of 95 °C for 10 minutes, followed by 40 cycles of 95 °C for 15 seconds and 60 °C for 60 seconds.

### 4.4. Tissue staining and imaging

Cecum tip harvested after euthanasia was immediately placed in 4 % PFA solution and fixed overnight. Cecum tip was then transferred to a 20 % sucrose solution for 4 hours and flash frozen in OCT embedding matrix (Sysmex Digitana, Switzerland). Embedded blocks were cut into 5 *μ*m sections and stored at *−*20 °C for FISH.

For FISH staining, slides were rinsed with PBS, followed by dehydration with increasing concentrations of ethanol. FISH probes were diluted to a final concentration of 1 %(v/v) in hybridization buffer and added to slides (Supplementary Tab. S5). Slides were incubated at 50 °C for 4 hours. Slides were washed with washing buffer at 50 °C for 20 minutes, then washed 3 more times with PBS at 50 °C for 20 minutes. Slides were air dried and mounted with Vectashield HardSet Antifade Mounting Medium (Vectashield). For imaging the EAM in the cecum, a Leica SP8 confocal microscope (Leica) was used at ScopeM (ETH Zürich).

### 4.5. Real-time Headspace-SESI measurements of bacterial cultures

The Headspace-SESI system consists of an in-house developed custom-made headspace sampling system (Fig 1C and Fig S1, adapted from [61]), a commercial SESI source (SuperSESI, Fossiliontech, Spain) and an Orbitrap mass spectrometer (Q-Exactive Plus, Thermo Scientific, San Jose, CA). All three EAM strains were able to grow in the Headspace-SESI system under the measurement condition (Fig S4A), including the strictly anaerobic bacteria *E. rectale*. A 20 μm Sharp Singularity Emitters (Fossiliontech, Spain) was used for electrospray using an aqueous solution of 0.1% formic acid. The sampling line temperature of the SESI source was set to 130 °C and the ion chamber temperature was set to 90 °C. To sample the bacterial culture’s headspace, 0.3 L min^*−*1^ of humidified carrier gas was flushed through a quartz culture flask containing 4 mL bacterial culture, and introduced into the SESI source. The humidifier, the headspace sampler and all gas tubing in between were kept at 37 °C. Optical density at wavelength = 595 nm was recorded in parallel. Detailed installation and light paths of the OD monitoring system were explained in Fig S1B

### 4.6. Real-time SESI measurements of mice

When measuring volatile released from mice, the headspace sampling system was replaced by an air-tight mouse tunnel (7 cm diameter, 11 cm length). The mouse tunnels were fabricated from poly(methyl methacrylate) (PMMA) and had polypropylen (PP) caps. The mouse tunnel was wrapped in red plastic foil, in order to reduce stress of the experiment animals. Nine 8-9 week old mixed sex germ-free, EAM-colonized, or SPF mice were housed under sterile conditions and kept on an *ad libitum* diet for 1 week. At the start of the experiment, mice were fasted from 6 am (zeitgeber 0). At zeitgeber 6, mice were restrained and urination was stimulated, and subsequently transferred individually to a mouse tunnel connected to SESI-MS. 0.3 L min^*−*1^ of humidified medical air (Pangas, Switzerland) was flushed through the mouse tunnel, and introduced into the SESI source. The humidifier, the mouse tunnel and all gas tubing in between were kept at room temperature (22 °C).

### 4.7. Mouse D-arabinose feeding

A visual summary of the D-arabinose feeding and measurement is in Fig 4A. Nine 8-9 week old mixed sex germ-free, EAM-colonized, or SPF mice were housed under sterile conditions and kept on an ad libitum diet for 1 week. At the start of the experiment, mice were fasted from 6 am (zeitgeber 0). At zeitgeber 3, 3 mice were given a 100 *μ*L mixture of 40 mg U-13C labeled D-arabinose and 40 mg unlabeled D-arabinose via oral gavage, with 10 minutes between each gavage. Five hours post-gavage, mouse headspace was measured in the SESI Orbitrap system for 500 seconds for each mouse.

Taking into account the fact that the orbitrap technology may suffer from deviating relative isotopic abundance caused by matrix[62], we measured mice without 13C-labeled arabinose as the baseline measurement to be compared with the mice at the same condition but fed with D-arabinose, so that their background metabolome are comparable. Mouse colonization, arabinose feeding and SESI-MS measurements were reproducible in a further experiment with less animals and in which a 6 hour time window was used between arabinose feeding and volatilome measurements (Fig S6)

### 4.8. MS settings for SESI measurements

Data was acquired on the Orbitrap in profile mode. For both positive and negative mode acquisition, the following settings were used: an ion transfer tube temperature 250 °C, a resolution of 140’000 at *m/z* = 200, RF lens set to 55%, and a maximum injection time of 100 ms. The automatic gain control (AGC) target was set to 5*·*10^6^. For bacterial volatiles, data were acquired with full scan over a range from *m/z* = 50-500. For mouse volatiles, data were acquired with a window-splitting method previous described by Lan et al. [63], where the whole mass range was covered by four narrow mass windows: for positive mode, *m/z* = 50-127, 127-143, 143-223, 223-500; for negative mode, *m/z* = 50-89, 89-197, 197-232, 232-500. Details for the MS acquisition method are shown in Fig S2.

### 4.9. Mass spectrometer data processing

Raw data generated from SESI-MS were first converted into .mzXML format via MSConvert (ProteoWizard [64]). The converted files were then imported into Matlab (version R2019b, MathWorks, Natick, MA) and treated using a Matlab code developed in-house. The following workflow is shown in Fig S3. Briefly, peaks were picked from an overall averaged spectrum of acquired spectra during the experiment. The acquired data was then centroided using the peak boundaries defined from the master spectra, and a .csv file containing the intensities of the peaks per scan was generated for each of the acquired MS data file.

### 4.10. Statistics

All statistics were done using in-house developed R codes. For prepossessing mass spectrometer data, the raw intensities were first normalized against total ion current, then the log2 of median intensities of the acquired scans were used as quantification. The pathway enrichment was done using the mummichog function of MetaboanalystR package [65, 66].

## Supporting information

Extended version of table S1

## Acknowledgments

We gratefully acknowledge financial support from SNF(No. 1851228) to G.G., from Heidi Ras Stiftung and Zurich Foundation to B.S. E.S. acknowledges the support of a Novartis Freenovation grant (2015), Swiss National Science Foundation Project grant (310030 185128), Botnar Research Center for Child Health MIP 2020 “Microbiota Engineering for Child Health” and the NCCR Microbiomes, a National Centre of Competence in Research, funded by the Swiss National Science Foundation (grant number 180575). We thank ScopeM (ETH Zurich) for providing us access to the confocal microscope, we thank RCHCI and EPIC for housing experiments animals. We also thank Anna Sintsova for helps with data treatment, Christoph Bärtschi and Benedikt Wanner for building the hardwares and electronics of the headspace sampling system. This work is part of the Zurich Exhalomics flagship project under the umbrella of “Hochschulmedizin Zürich”.

## Author contributions

J.L., G.G., M.A., R.Z. and E.S. designed the research. J.L. performed the mass spectrometry experiments and data evaluation, with technical advice from B.S. G.G. performed the biological experiments and data evaluation. J.L., M.A. and E.S. wrote the manuscript, to which G.G. contributed, in cooperation with all other authors. R.Z. and E.S. acquired funding supports.

