## Supplementary Figures 1-7 for "Non-invasive monitoring of microbiota and host metabolism using Secondary electrospray ionization-Mass spectrometry"

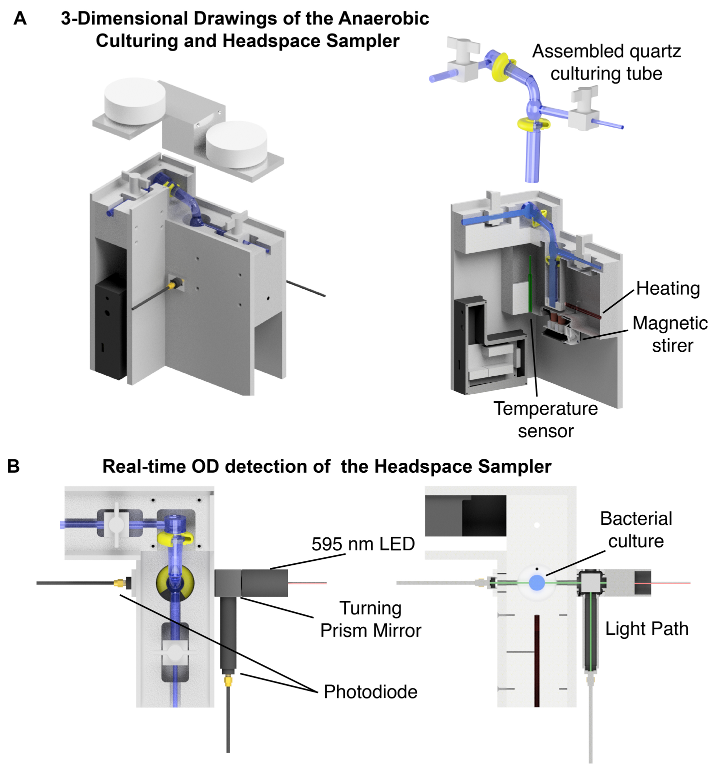


Figure S1: **Detailed design of anaerobic culturing and headspace sampler. A,** 3-dimensional drawing of the heating block and an assembled quartz culturing tube. **B,** real time optical density detector of the headspace sample.

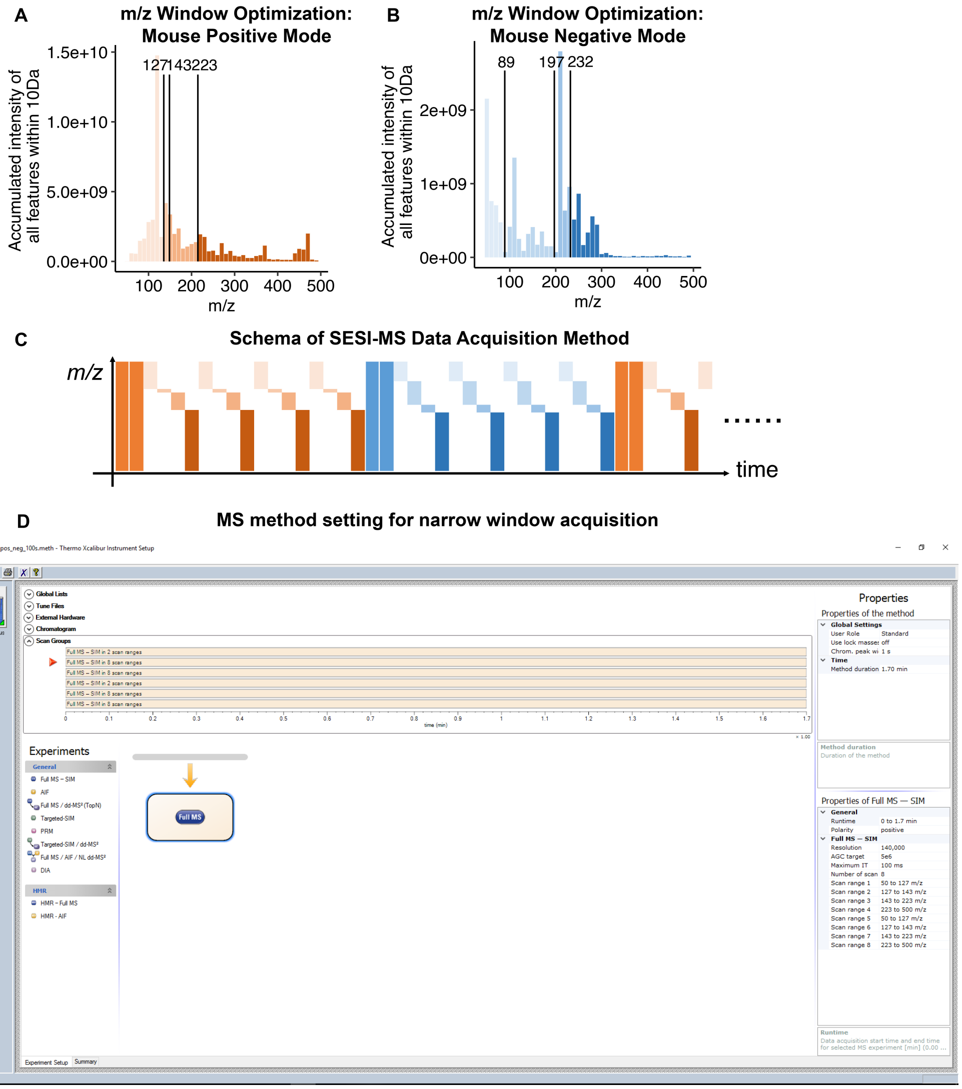


Figure S2: **Narrow windows acquisition method for SESI-MS. A,** Positive and **B,** negative polarity ion distribution of mouse volatiles, measured by twenty-two stitched 20 Da windows. Mass windows were defined to ensure an equal cumulative intensity per window. **C,** Schema of SESI-MS data acquisition method using narrow mass windows. Briefly, for each polarity, two full scans were first measured, followed by scanning through $m/z = 50$-500 using narrow mass windows four times. **D,** MS method setting for narrow mass windows acquisition, demonstrated in Xcalibur software (Thermo Scientific).

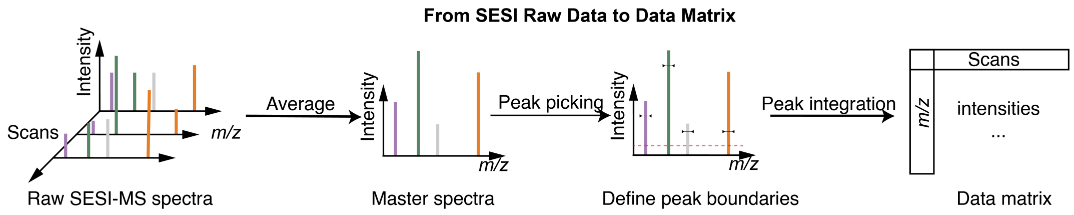


Figure S3: Pipeline for generated data matrix from SESI raw data. (See the method section 4.9 for details.)

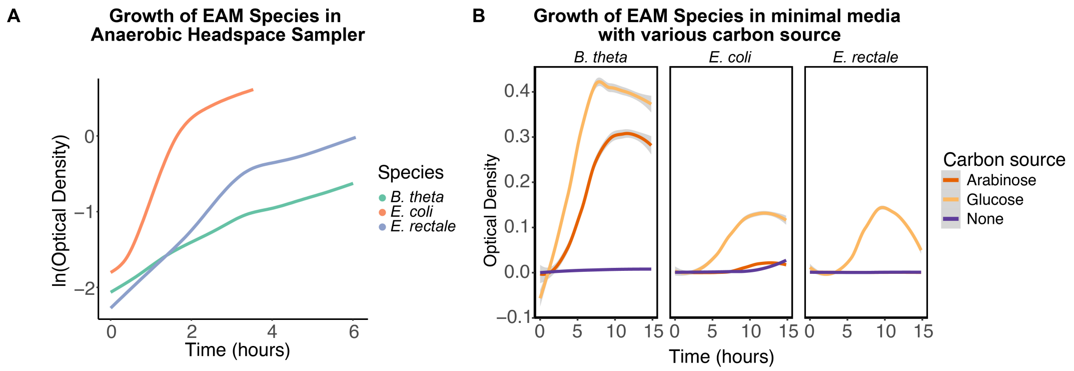


Figure S4: **Growth of EAM species under different conditions A,** in anaerobic headspace sampler coupling to SESI-MS, in BHIS. **B,** in anaerobic tent with minimal media supplied with different carbon sources.

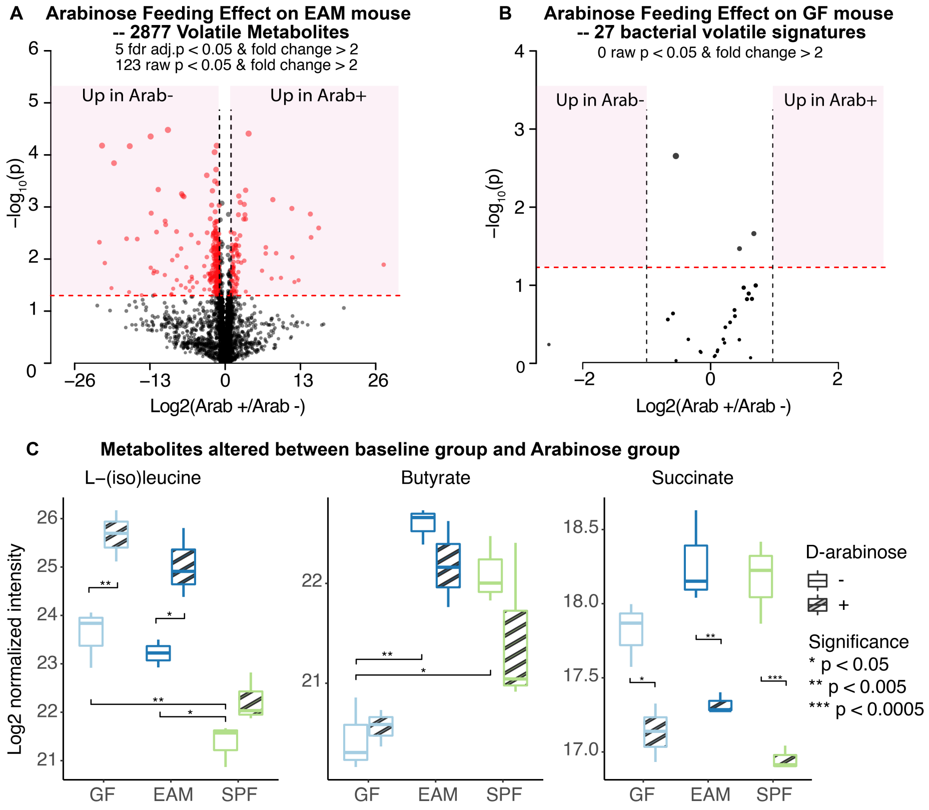


Figure S5: **Altered total metabolome between D-arabinose +/- EAM mouse. A,** Volcano plot showing metabolites altered between baseline group and arabinose group. (5 FDR adj. $p < 0.05$ & fold change $> 2$, 123 raw $p < 0.05$ & fold change $> 2$) **B,** Volcano plot showing arabinose feeding effect on bacterial volatile signatures detected in GF mice. (0 raw $p < 0.05$ & fold change $> 2$) **C,** Examples of altered volatile metabolites detected by SESI.

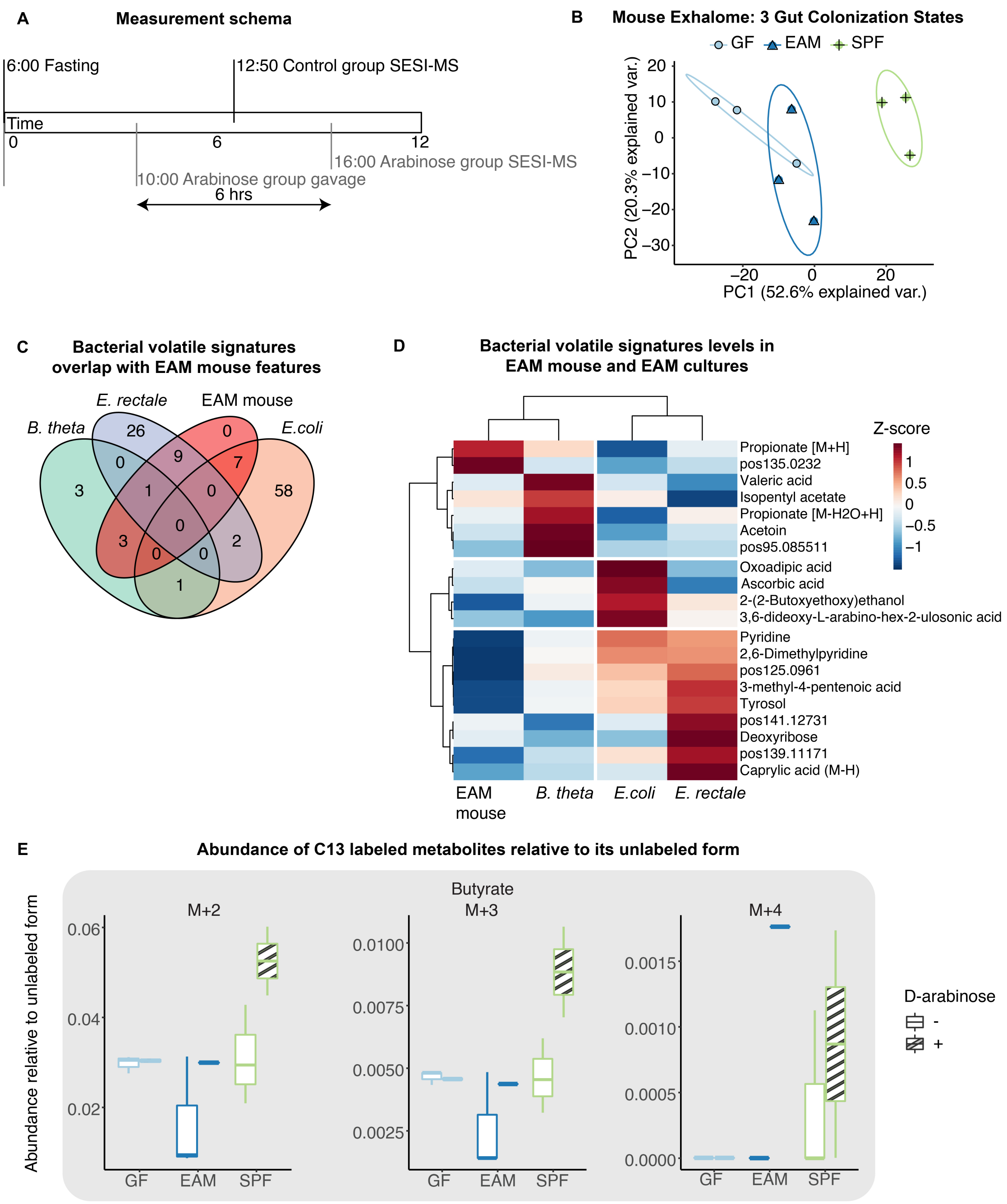


Figure S6: **Independent technical and biological replicates show the Key findings are reproducible in an experiment performed six months apart. A,** Measurement schema of the control group and the arabinose group. **B,** PCA plot of volatilome of mice with different gut microbiota: germ-free, EAM-colonized and SPF. **C,** Overlap between bacterial volatile signatures identified *in vitro* and EAM mouse volatile features. **D,** Hierarchical clustering showing normalized intensities of bacterial volatile signatures in culture and in EAM-colonized mouse. **E,** Labelling patterns of butyrate released by mice with different microbiota.

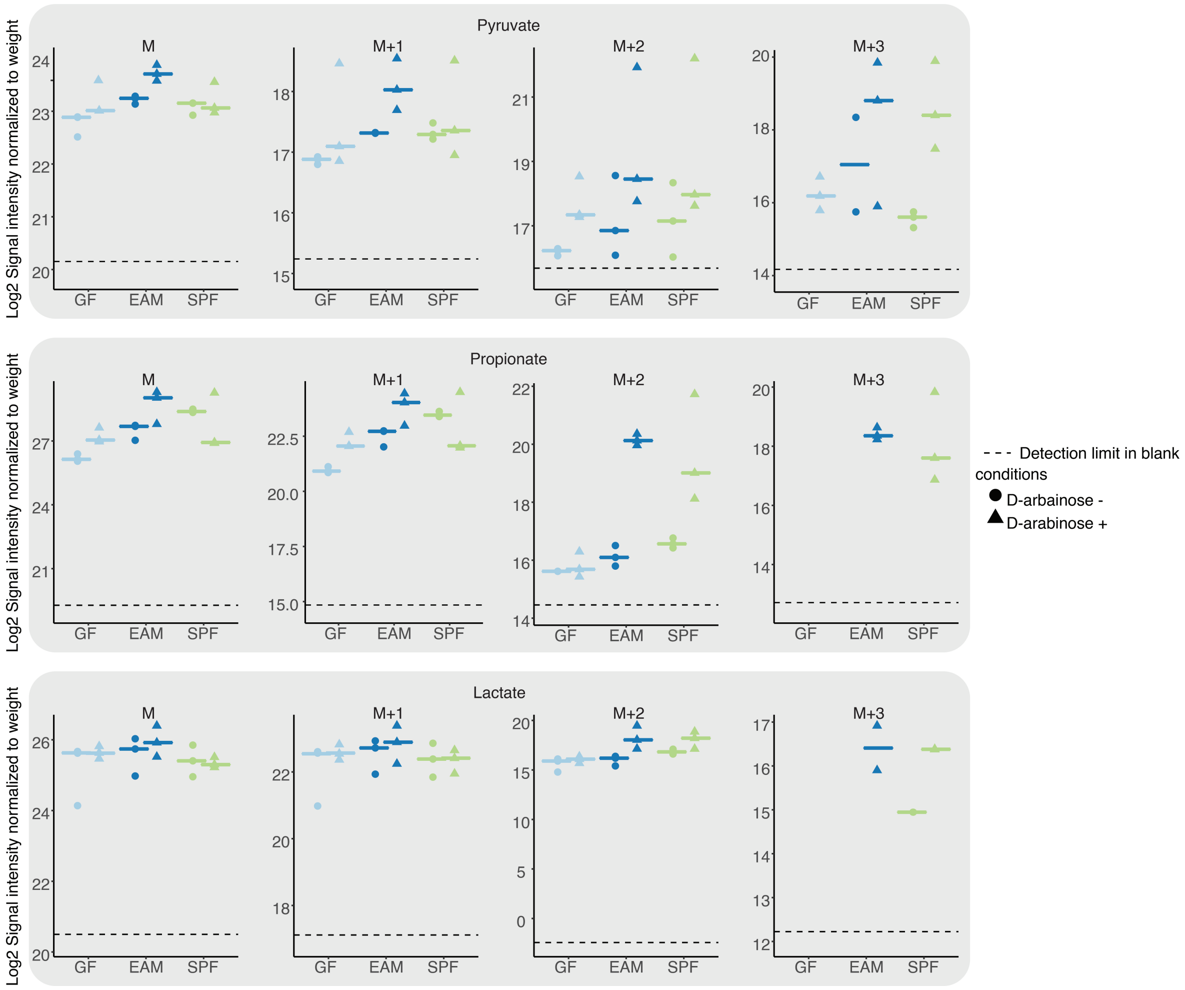


(continue on the next page)

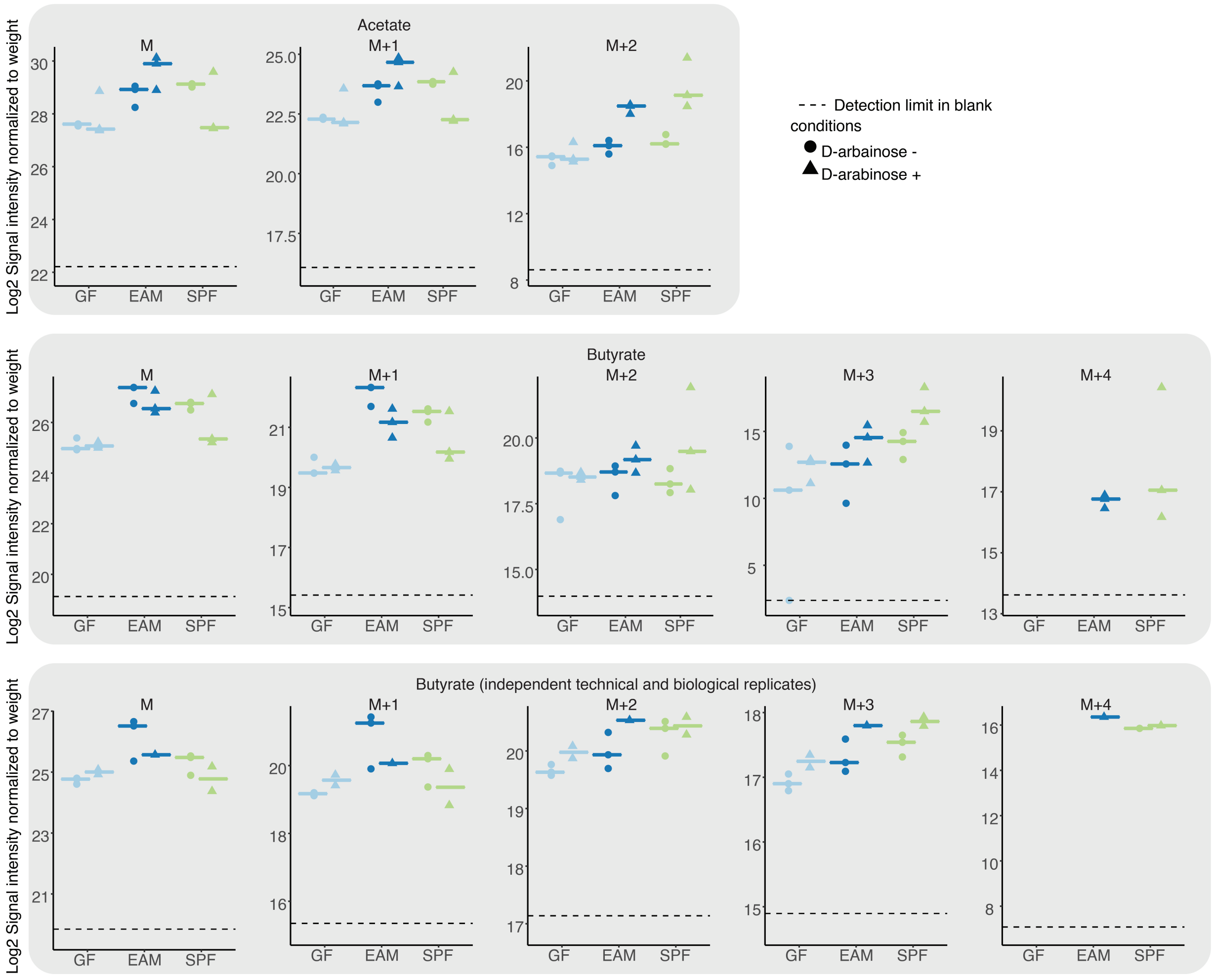


Figure S7: **Log2 intensity of the C13 labeled metabolites**
