## Supplementary tables 1-5 for "Non-invasive monitoring of microbiota and host metabolism using Secondary electrospray ionization-Mass spectrometry"

Table S1: Bacterial volatile signatures identified *in vitro*. The numbers in the "source" column indicate the following annotation methods: 1-described in a previous study as bacterial volatile metabolites[67] 2-in KEGG database, 3-MS2 matching with METLIN database[42], 4-MS2 matching with MassBank [43] using MSDIAL software [68], 5-formula only and 0-No available annotation

| strain | feature | Ionization form/Formula | Final annotation | source |
|---|---|---|---|---|
| medium | pos139.07531 | M+H | tyrosol | 2 |
| medium | neg125.02427 | M-H | Hydroxyhydroquinone | 2,3 |
| medium | pos115.07531 | M+H | 3-methyl-4-pentenoic acid | 3 |
| medium | pos101.05978 | M-NH3+H | L-Valine | 2 |
| medium | neg109.02761 | M-H | Catechol | 2,3,4 |
| medium | neg115.03474 | NA | NA | 0 |
| medium | neg205.15941 | M-H | 4-Octylphenol | 2,3 |
| B. theta | pos95.085511 | C7H11+ | C7H11+ | 5 |
| B. theta | pos71.04915 | M-H2O+H | Acetoin | 1,2 |
| B. theta | pos103.07537 | M+H | Valeric acid | 1,3 |
| B. theta | neg101.05739 | M-H | Valeric acid | 1,3 |
| B. theta | neg114.03116 | M-H2O-H | (2S)-amino(carbamoylamino)ethanoic acid | 2 |
| B. theta | neg115.03898 | C3H5N3O2- | C3H5N3O2- | 5 |
| B. theta | pos113.09598 | C7H13O + | C7H13O + | 5 |
| B. theta | pos131.10658 | M+H | Isopentyl acetate | 1 |
| E. rectale | neg101.021 | NA | NA | 0 |
| E. rectale | neg103.03662 | M(C13)-H | hydroxy butyric acid | 2,3,4 |
| E. rectale | neg107.0475 | M-H | p-Cresol | 3 |
| E. rectale | neg111.04324 | C4H5N3O - | C4H5N3O - | 5 |
| E. rectale | neg115.07539 | M-H | Caproic acid | 1 |
| E. rectale | neg131.07121 | M-H | 2-Ethyl-2-Hydroxybutyric acid | 3 |
| E. rectale | neg133.04871 | M(C13)-H | Mandelonitrile | 2 |
| E. rectale | neg133.0505 | M-H | Deoxyribose | 2 |
| E. rectale | neg135.06618 | M+FA-H | meso-2,3-Butanediol | 2 |
| E. rectale | neg143.10758 | M-H | Caprylic acid | 2,3 |
| E. rectale | neg144.11089 | M(C13)-H | Isotope | 2,3 |
| E. rectale | neg149.04533 | M-H | Ribose | 2 |
| E. rectale | neg205.15941 | M-H | 4-Octylphenol | 3 |
| E. rectale | neg206.16286 | M(C13)-H | Isotope | 3 |
| E. rectale | neg251.16494 | M+FA-H | 4-Octylphenol | 3 |
| E. rectale | neg87.041311 | NA | NA | 0 |
| E. rectale | neg88.044651 | NA | Isotope | 0 |
| E. rectale | neg99.041011 | NA | NA | 0 |
| E. rectale | pos108.08075 | M+H | 2,6-Dimethylpyridine | 3 |
| E. rectale | pos115.07531 | M+H | 3-methyl-4-pentenoic acid | 3 |
| E. rectale | pos118.06506 | M+H | indole | 1,2,3 |
| E. rectale | pos119.05248 | C5H11OS + | C5H11OS + | 5 |
| E. rectale | pos122.09642 | M+H | 2,6-Xylidine | 3 |
| E. rectale | pos125.0961 | C8H13O + | C8H13O + | 5 |
| E. rectale | pos129.01578 | M+Na | glyceric acid | 2 |
| E. rectale | pos131.10658 | M+H | Pentyl acetate | 1,2 |
| E. rectale | pos133.0859 | M-NH3+H | Triethanolamine | 2 |
| E. rectale | pos135.08038 | M+H | 2,4-Dimethylbenzaldehyde | 3 |
| E. rectale | pos139.07531 | M+H | tyrosol | 2 |
| E. rectale | pos139.11171 | C9H15O + | C9H15O + | 5 |
| E. rectale | pos141.12731 | C9H17O+ | C9H17O+ | 5 |
| E. rectale | pos147.10149 | M+H2O+H | Cyclohexanecarboxylic acid | 2 |
| E. rectale | pos149.09602 | M+H-H2O | Perillic acid | 2 |
| E. rectale | pos171.13788 | M+H-H2O | Citronellic acid | 3 |
| E. rectale | pos182.90232 | NA | NA | 0 |
| E. rectale | pos197.15353 | M+H | pentamethylmelamine | 3 |
| E. rectale | pos209.15352 | C13H21O2+ | C13H21O2+ | 5 |
| E. rectale | pos80.049511 | M+H | Pyridine | 4 |
| E. coli | neg108.04274 | C3H8O4 - | C3H8O4 - | 5 |
| E. coli | neg109.02761 | M-H | Catechol | 3,4 |
| E. coli | neg110.02284 | C3H2N4O - | 2,6-Dihydroxypyridine | 3 |
| E. coli | neg112.00211 | C2N4O2 - | C2N4O2 - | 5 |
| E. coli | neg112.0385 | C3H4N4O - | C3H4N4O - | 5 |
| E. coli | neg112.99734 | NA | NA | 0 |
| E. coli | neg114.01861 | M-H2O-H | L-Aspartic acid | 2,3 |
| E. coli | neg114.03116 | M-H2O-H | (2S)-amino(carbamoylamino)ethanoic acid | 2 |

| strain | feature | Ionization form/Formulae | Final annotation | source |
|---|---|---|---|---|
| E. coli | neg116.00863 | NA | NA | 0 |
| E. coli | neg116.01043 | C2H2N3O3 - | C2H2N3O3 - | 5 |
| E. coli | neg116.09044 | NA | NA | 0 |
| E. coli | neg117.04658 | C7H5N2 - | C7H5N2 - | 5 |
| E. coli | neg117.0529 | CH5N6O - | hydroxyvaleric acid | 4 |
| E. coli | neg117.0546 | M+FA-H | Butanal | 2 |
| E. coli | neg118.05624 | M(C13)-H-H2O | isotope | 2 |
| E. coli | neg120.04538 | M-H | Benzamide | 2 |
| E. coli | neg122.02466 | M-H | Nicotinic acid | 2 |
| E. coli | neg124.04024 | C6H6NO2 - | C6H6NO2 - | 5 |
| E. coli | neg130.01442 | M-H | 2-Oxosuccinamate | 2 |
| E. coli | neg132.04533 | M-H | Mandelonitrile | 2 |
| E. coli | neg135.06618 | M+FA-H | 1,3-Butanediol | 2 |
| E. coli | neg136.04023 | M-H | 2-Aminobenzoic acid(C00108) | 2,3,4 |
| E. coli | neg137.02427 | M-H | 4-Hydroxybenzoic acid(C00156) | 2,3,4 |
| E. coli | neg138.01951 | M-H | 4-Nitrophenol (C00870) | 2,3,4 |
| E. coli | neg139.03993 | M+FA-H | Phenylic alcohol(C00146) | 2 |
| E. coli | neg141.01918 | M-H-H2O | Oxoadipic acid | 2 |
| E. coli | neg143.017 | M-H-H2O | 1,2-Dihydroxy-3-keto-5-methylthiopentene | 2 |
| E. coli | neg143.03478 | M-H2O-H | 3,6-dideoxy-L-arabino-hex-2-ulosonic acid(C03979) | 2 |
| E. coli | neg146.02461 | M-H | Indole-5,6-quinone | 2 |
| E. coli | neg147.03239 | C8H5NO2 - | C8H5NO2 - | 5 |
| E. coli | neg148.04028 | M-H2O-H | Pyridoxal | 2 |
| E. coli | neg149.04797 | C5H10ClN2O - | C5H10ClN2O - | 5 |
| E. coli | neg154.01439 | M-H | 4-Nitrocatechol | 2 |
| E. coli | neg154.05081 | M-H2O-H | N-Acetyl-L-glutamate 5-semialdehyde(C01250) | 2 |
| E. coli | neg155.03481 | M-H2O-H | Shikimic acid | 1,2 |
| E. coli | neg157.01402 | M-H2O-H | Ascorbic acid | 2 |
| E. coli | neg159.02968 | M-H | Oxoadipic acid | 2,3 |
| E. coli | neg163.02729 | C8H5NO3 - | C8H5NO3 - | 5 |
| E. coli | neg165.03842 | M(C13)-H | 4-Pyridoxolactone | 2 |
| E. coli | neg165.04288 | M(C13)-H | (2S)-2-amino-4-(methylsulfinyl)butanoic acid(C02989) | 2 |
| E. coli | neg166.05072 | M-H | Pyridoxal (Vitamin B6) | 2,3,4 |
| E. coli | neg168.02999 | M+FA-H | Nicotinic acid | 2 |
| E. coli | neg168.0664 | M-H | Pyridoxine(C00314) | 2,4 |
| E. coli | neg171.0297 | M-H2O-H | 3-Dehydroquinate | 2 |
| E. coli | neg172.02499 | C6H6NO5- | C6H6NO5- | 5 |
| E. coli | neg180.03005 | M-H | 2-Methyl-3-hydroxy-5-formylpyridine-4-carboxylate | 2 |
| E. coli | neg181.03786 | NA | NA | 0 |
| E. coli | neg182.04553 | M+FA-H | 4-pyridoxate | 4 |
| E. coli | neg196.02482 | M-H | 3-Hydroxy-2-methylpyridine-4,5-dicarboxylate | 2 |
| E. coli | neg198.0406 | M+FA-H | 3-Hydroxyanthranilic acid(C00632) | 2 |
| E. coli | neg200.05624 | M-H2O-H | O-Succinylhomoserine(C01118) | 2 |
| E. coli | neg212.05626 | M+FA-H | Pyridoxal | 2 |
| E. coli | neg214.03546 | C8H8NO6- | C8H8NO6- | 2 |
| E. coli | neg246.06167 | M-H | N-(3-carboxypropanoyl)-L-glutamic acid(C05931) | 2 |
| E. coli | neg251.16494 | C15H23O3 - | C15H23O3 - | 5 |
| E. coli | neg59.98763 | NA | NA | 0 |
| E. coli | neg60.99476 | NA | NA | 0 |
| E. coli | neg60.99818 | NA | NA | 0 |
| E. coli | neg85.025741 | NA | NA | 0 |
| E. coli | neg97.025421 | NA | NA | 0 |
| E. coli | pos102.12776 | C6H16N+ | C6H16N+ | 5 |
| E. coli | pos119.06849 | M(C13)+H | Indole | 2,3 |
| E. coli | pos135.0232 | NA | NA | 0 |
| E. coli | pos147.08035 | M-H2O+H | 2-Phenylethyl acetate | 1 |
| E. coli | pos163.13278 | M+H2O+H | 2-(2-Butoxyethoxy)ethanol | 3 |
| E. coli | pos57.03356 | M-H2O+H | Propionate | 2 |
| E. coli | pos75.044051 | M+H | Propionate | 2 |
| E. coli | pos97.026061 | M+Na | Propionate | 2 |

Table S2: EAM colonization effect on host metabolism: pathway enrichment.

| KEGG pathways | Enrichment factor | Fisher's exact test, p value |
|---|---|---|
| Butanoate metabolism | 6.837 | 0.047 |
| Aminoacyl-tRNA biosynthesis | 4.662 | 0.140 |
| Glyoxylate and dicarboxylate metabolism | 3.309 | 0.181 |
| Propanoate metabolism | 4.048 | 0.184 |
| Caffeine metabolism | 4.273 | 0.096 |
| Cysteine and methionine metabolism | 2.331 | 0.357 |
| Drug metabolism - cytochrome P450 | 2.442 | 0.246 |
| Arginine and proline metabolism | 3.465 | 0.506 |
| Glycine, serine and threonine metabolism | 3.309 | 0.515 |
| Pyrimidine metabolism | 1.972 | 0.472 |

Table S3: Arabinose feeding effect on EAM mouse metabolism: pathway enrichment.

| KEGG pathways | Enrichment factor | Fisher's exact test, p value |
|---|---|---|
| Valine, leucine and isoleucine degradation | 4.579 | 0.002 |
| Butanoate metabolism | 7.478 | 0.001 |
| Pentose and glucuronate interconversions | 4.713 | 0.011 |
| Phenylalanine metabolism | 5.342 | 0.009 |
| Sphingolipid metabolism | 7.122 | 0.009 |
| Biosynthesis of unsaturated fatty acids | 1.885 | 0.009 |
| Valine, leucine and isoleucine biosynthesis | 10.016 | 0.061 |
| Alanine, aspartate and glutamate metabolism | 4.006 | 0.099 |
| Citrate cycle (TCA cycle) | 5.008 | 0.105 |

Table S4: Primer sequences for qPCR analysis of EAM species.

| Species | Primer | Primer sequence |
|---|---|---|
| *B. theta* | Forward | TACTCGCCTCTTTGCAACCCTACC |
| | Reverse | GGCCCCAGATCCGAACAACAC |
| *E. coli* | Forward | GGTGGCTGGGTGATGTAAAACTGA |
| | Reverse | ACCGCCGAGCAAAATGAAGC |
| *E. rectale* | Forward | GGTACAACAGGCGTTATTGTATC |
| | Reverse | CGAAAGCACCGATCTTCTT |

Table S5: Probes for *in vivo* EAM FISH imaging.

| FISH probes | Dye | Modification at | Sequence |
|---|---|---|---|
| *E. rectale* | Texas Red | 5’ | GCTTCTTAGTCAGGTACCG |
| *E. coli* | ATTO425 | 5’ | GCCTTCCCACATCGTTT |
| *B. theta 1* | Cy5 | 5’ | CCAATGTGGGGGACCTT [69] |
| *B. theta 2* | Cy5 | 5’ | CATTTGCCTTGCGGCTA [70] |
| *B. theta 3* | Cy5 | 5’ | AGCTGCCTTCGCAATCGG [71] |
